## Supplemental Figures and Datasets for "Opposite polarity programs regulate asymmetric subsidiary cell divisions in grasses"

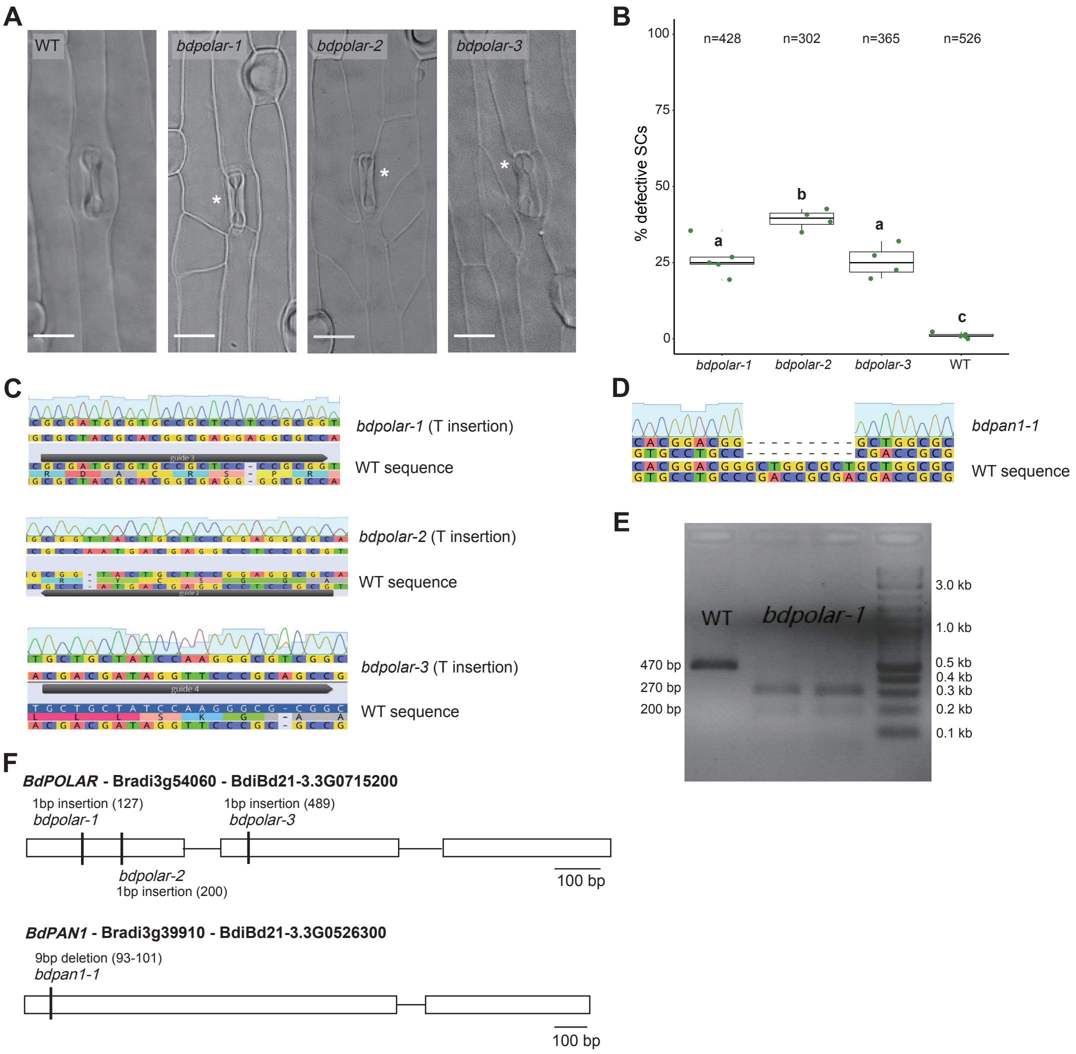


**Figure S1. CRISPR/Cas9 and EMS-mutagenized mutants in *BdPOLAR* and *BdPAN1*. (A)** DIC images of the epidermis in WT, *bdpolar-1*, *bdpolar-2,* and *bdpolar-3* (third leaf, 19 dag). Aberrant SCs are indicated with white asterisks. Scale bar, 15 µm. **(B)** Quantifications of defective SCs in *bdpolar-1*, *bdpolar-2*, *bdpolar-3,* and WT control. Samples were compared using a one-way ANOVA and post-hoc Tukey test for multiple comparisons; different letters indicate significant differences ($p<0.05$); n=4-5 individuals and 302-526 SCs. **(C)** Genotyping chromatogram of CRISPR/Cas9-induced mutations in *bdpolar-1*, *bdpolar-2,* and *bdpolar-3.* WT sequence at the bottom and mutant sequence chromatogram above are displayed; CRISPR/Cas9 guides are indicated. **(D)** Genotyping chromatogram of EMS-induced mutation in *bdpan1-1.* WT sequence at the bottom and mutant sequence chromatogram above are displayed. **(E)** Agarose gel electrophoresis of the BseRI-digested CAPS marker to genotype *bdpolar-1*. BseRI can digest PCR products from *bdpolar-1* (200bp, and 270bp) but not from WT (470bp). DNA kilobases standard control is NEB 1kb Plus DNA Ladder. **(F)** Gene model of *BdPOLAR* (Bradi3g54060, BdiBd21-3.3G0715200) with available CRISPR/Cas9-induced (*bdpolar-1*, *bdpolar-2*, *bdpolar-3*) mutations and *BdPAN1* (Bradi3g39910, BdiBd21-3.3G0526300) with EMS-induced mutation *bdpan1-1*.

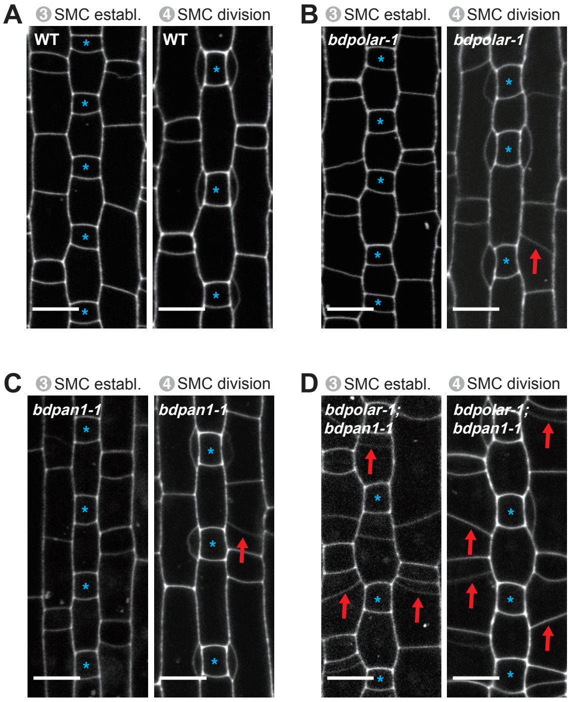


**Figure S2. Misoriented SMC division planes likely cause aberrant SCs in mature leaf zones.** Single confocal plane images of the PI-stained developing epidermis showing stage 3 to stage 4 SMCs in WT, *bdpolar-1*, *bdpan1-1*, and *bdpolar-1;bdpan1-1*. GMCs are indicated with blue asterisks. Red arrows indicate misoriented division planes in SMCs. Scale bar, 10 µm.

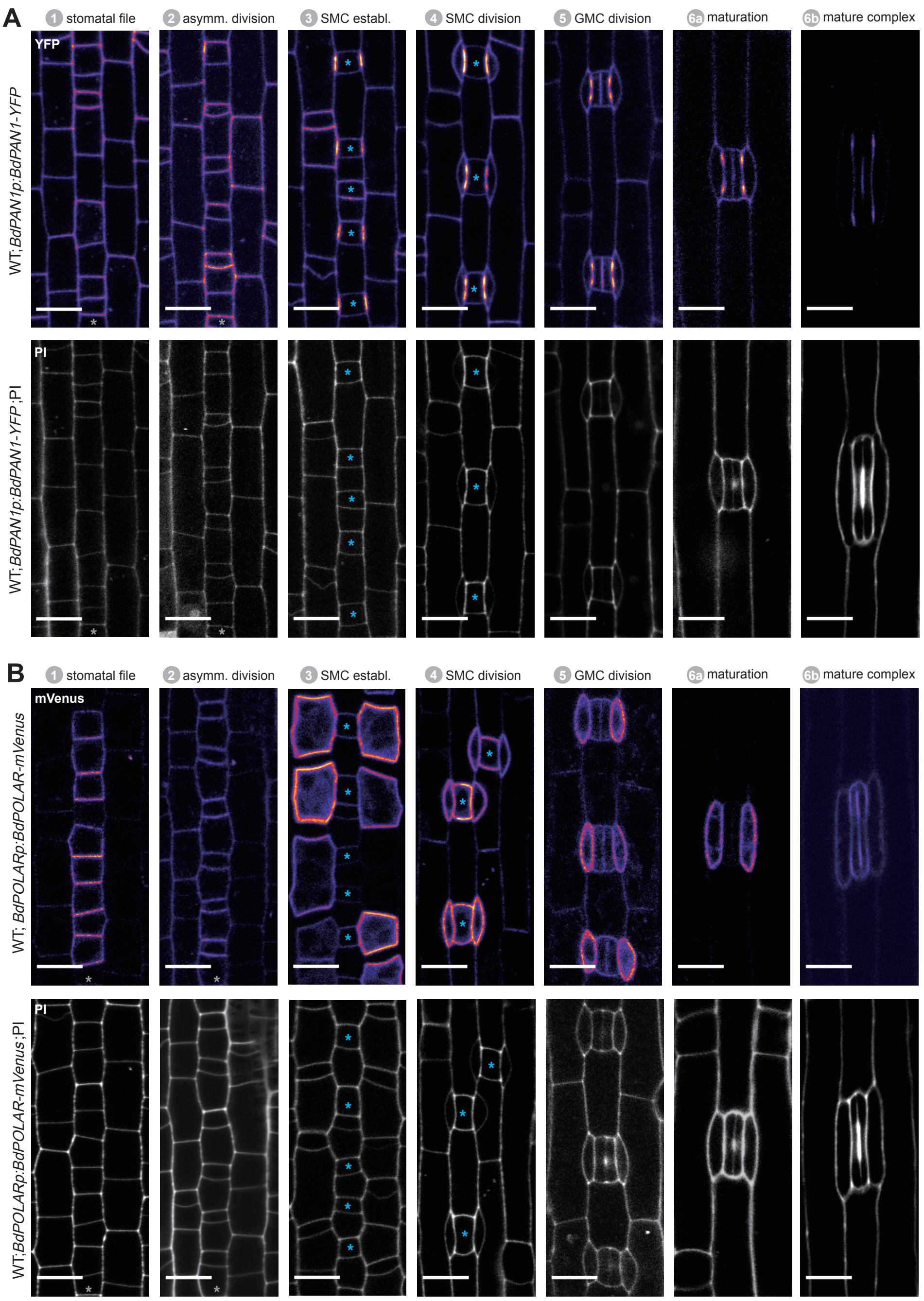


**Figure S3. BdPAN1 and BdPOLAR expression throughout stomatal development in *B. distachyon.* (A)** *BdPAN1p:BdPAN1-YFP* reporter expression during stomatal development. Fluorescence intensity images of YFP channel only (upper) and images of PI-stained cell outlines only (bottom). **(B)** *BdPOLARp:BdPOLAR-mVenus* reporter expression during stomatal development. Fluorescence intensity images of mVenus channel only (upper) and images of PI-stained cell outlines only (bottom). Note that the laser was adjusted for different stages to visualise the signal and expression level between stages cannot be quantitatively compared. The stomatal files are indicated with grey asterisks. GMCs are indicated with blue asterisks. Stomatal stages are indicated. Scale bar, 10 µm.

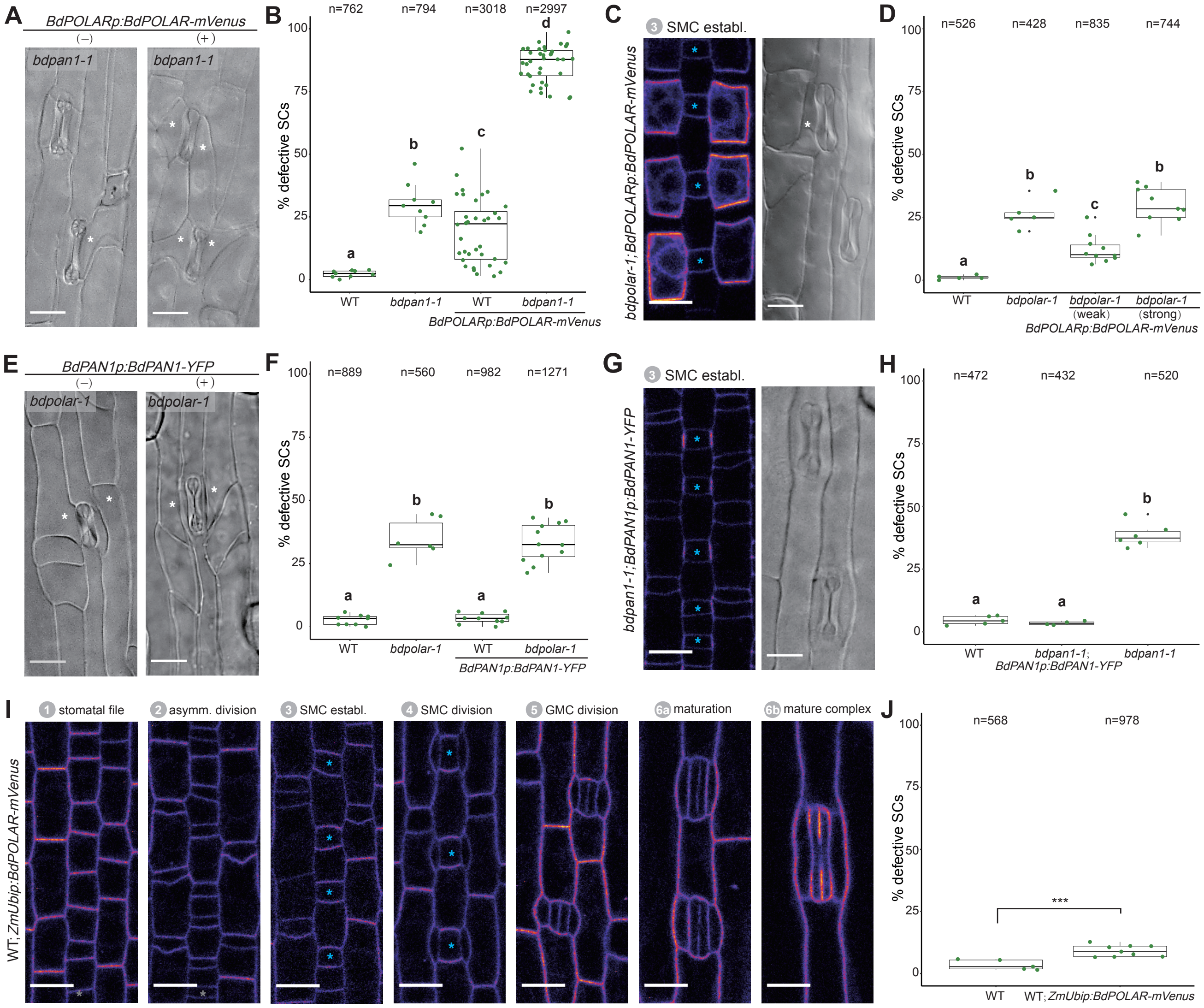


**Figure S4. Dosage of BdPOLAR is crucial for its function. (A)** DIC images of the epidermis in *bdpan1-1* with (+) or without (-) *BdPOLARp:BdPOLAR-mVenus* expression. **(B)** Quantifications of defective SCs in WT and *bdpan1-1* with or without *BdPOLARp:BdPOLAR-mVenus* expression; n=9-38 individuals and 762-3018 SCs. **(C)** Fluorescence intensity image of *BdPOLARp:BdPOLAR-mVenus* in *bdpolar-1,* and DIC images (right) of the mature epidermis of *bdpolar-1* complemented with *BdPOLARp:BdPOLAR-mVenus*. **(D)** Quantifications of defective SCs in WT, *bdpolar-1* complemented with weakly and strongly expressed *BdPOLARp:BdPOLAR-mVenus*, and *bdpolar-1* control. n=5-10 individuals and 428-835 SCs. **(E)** DIC images of the epidermis in *bdpolar-1* with (+) or without (-) *BdPAN1p:BdPAN1-YFP* expression. **(F)** Quantifications of defective SCs in WT and *bdpolar-1* with or without *BdPAN1p:BdPAN1-YFP* expression; n=6-12 individuals and 560-1271 SCs. **(G)** Fluorescence intensity image of *BdPAN1p:BdPAN1-YFP* in *bdpan1-1* (left), and DIC images of the epidermis in *bdpan1-1* complemented with *BdPAN1p:BdPAN1-YFP* (right). **(H)** Quantifications of defective SCs of WT, *bdpan1-1* complemented with *BdPAN1p:BdPAN1-YFP*, and *bdpan1-1* control. Data for WT is the same as in Fig. S4D and data for *bdpan1-1* control is the same as in Fig. 1D. n=4-6 individuals and 432-520 SCs. **(I)** Fluorescence intensity image for *ZmUbip:BdPOLAR-mVenus* reporter expression during stomatal development. Stages are indicated. **(J)** Quantifications of defective SCs of WT and WT with *ZmUbip:BdPOLAR-mVenus* expression; n=5-9 individuals and 568-978 SCs. Samples were compared using a one-way ANOVA and posthoc Tukey test for multiple comparisons; different letters indicate significant differences ($p<0.05$). For comparisons between two groups, a Welch t-test was used; *** = p-value < 0.001. Aberrant SCs are indicated with white asterisks. The stomatal files are indicated with grey asterisks. GMCs are indicated with blue asterisks. Scale bar in DIC images, 15 $\mu$m; Scale bar in confocal images, 10 $\mu$m.

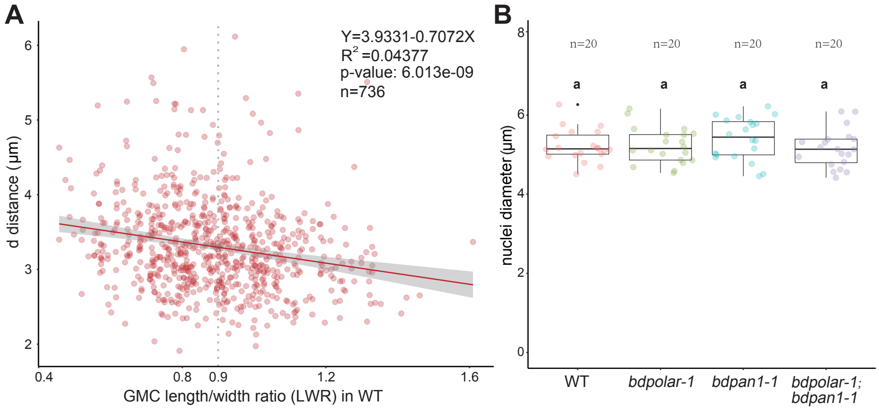


**C** GMC LWR > 0.9

| Genotypes | Total nuclei (n) | d distance (µm) | Polarized nuclei (n) | Nuclei polarization % |
|---|---|---|---|---|
| WT | 339 | 3.189794 | 316 | 93.22% |
| *bdpolar-1* | 590 | 3.476963 | 469 | 79.49% |
| *bdpan1-1* | 376 | 3.775290 | 243 | 64.63% |
| *bdpolar-1;bdpan1-1* | 445 | 3.888517 | 279 | 62.70% |

**Figure S5. *BdPAN1* controls nuclear polarisation. (A)** Scatter plot displaying d distance (*µ*m; y-axis) as a function of GMC length/width ratio (LWR; x-axis) in SMCs of WT (n=736 SMCs). Gray dashed line indicates GMC LWR=0.9. The linear regression is indicated and the regression equation, $R^2$ value, and p-value are indicated at the top right. **(B)** Quantification of the SMC nuclei diameter in WT, *bdpolar-1*, *bdpan1-1* and *bdpolar-1*;*bdpan1-1* (n=20 SMCs). Samples were compared using a one-way ANOVA and post-hoc Tukey test for multiple comparisons; different letters indicate significant differences (p<0.05). **(C)** Summary table of total number of quantified nuclei, average distance (d [*µ*m]) when GMC LWR>0.9, and number and percentage of nuclei within 4 *µ*m range when GMC LWR>0.9 (=polarised nuclei) in WT, *bdpolar-1*, *bdpan1-1* and *bdpolar-1*;*bdpan1-1*.

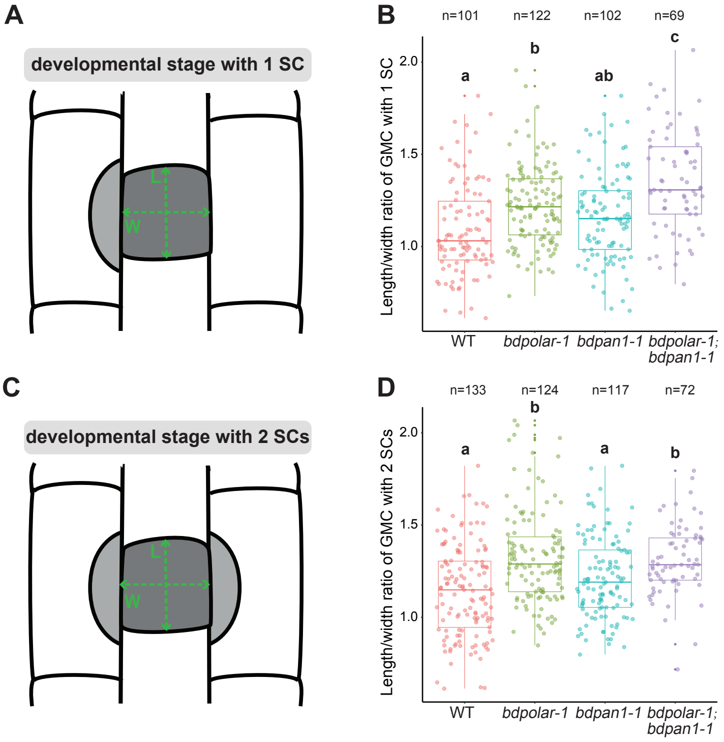


**Figure S6. GMCs that successfully recruited SCs displayed a higher GMC length to width ratio (LWR) in *bdpolar-1* compared to *bdpan1-1.*** **(A, C)** Schematic showing GMCs that recruited one (A) or two SCs (C). L: GMC length; W: GMC width. **(B, D)** Quantifications of GMC LWR in WT, *bdpolar-1*, *bdpan1-1* and *bdpolar-1*;*bdpan1-1* when GMCs recruited one (B) and two SCs (D); Samples were compared using a one-way ANOVA and post-hoc Tukey test for multiple comparisons; different letters indicate significant differences ($p<0.05$). Numbers of GMCs analysed are indicated.

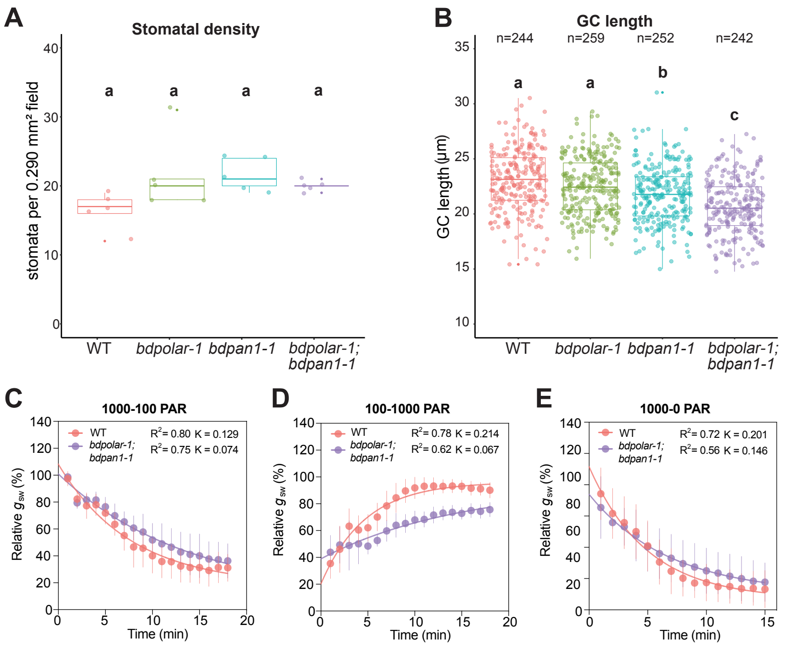


**Figure S7. The stomatal density and GC length in WT, *bdpolar-1, bdpan1-1* and *bdpolar-1;bdpan1-1*. (A)** Stomatal density was quantified using the leaf areas that were used for stomatal conductance measurements in Fig. 5. n=5 individuals and 364 - 418 stomatal complexes. **(B)** GC length was quantified using the leaf areas that were used for stomatal conductance measurements in Fig. 5. n=5 individuals and 242 - 259 GCs. **(C)** One-phase decay exponential regression for the transition 1000 to 100 PAR (n = 5 individuals). **(D)** One-phase association exponential regression for the transition 100 to 1000 PAR (n = 5 individuals). **(E)** One-phase decay exponential regression for the transition 1000 to 0 PAR (n = 5 individuals). $R^2$ value and constant rate (K) are indicated. Samples were compared using a one-way ANOVA and post-hoc Tukey test for multiple comparisons; different letters indicate significant differences (p<0.05).

**SUPPLEMENTAL TABLES, DATASETS AND VIDEOS**

**Table S1. Differentially expressed genes in bdmute developing leaf zones; related to Fig. 1.**

**Table S2. Primers used in this study.**

**Table S3. Stomatal conductance data in WT, *bdpolar-1, bdpan1-1* and *bdpolar-1;bdpan1-1*.**

**Supplementary dataset 1. Quantification data reported in this paper.**

**Video S1. Animated 3D rendering of BdPAN1-YFP domain in SMCs.**

**Video S2. Animated 3D rendering of BdPOLAR-mVenus domain**
